# ceQTL: A co-expression QTL model to detect a variant that affects transcription factor binding and its target regulation

**DOI:** 10.1101/2025.10.06.680553

**Authors:** Panwen Wang, Li Liu, Junwen Wang, Ping Yang, Zhifu Sun

## Abstract

Expression quantitative trait locus (eQTL) mapping is used to identify a functional link between a genomic variant, such as single nucleotide polymorphism (SNP), and gene expression (often close-by pair for cis-eQTL) by linear regression, commonly done when matching genotype and expression data from same individuals are available. Millions of significant eQTLs have been reported by both individual studies and coordinated large consortiums such as Genotype-Tissue Expression project (GTEx). A significant eQTL association does not establish a causal relationship or provide any underlying mechanism so further investigation is needed to understand how a SNP impacts gene expression. One of the plausible explanations for eQTL is that a genomic variant affects transcription factor (TF) binding and thus impacts its regulation on its target genes (TGs). However, data-driven or formal statistical methods to prove that hypothesis are still lacking. To address the gap, we propose a new method called differential co-expression QTL (ceQTL) among different alleles using Chow statistics to specifically detect eQTLs that are modulated by a particular TF. We start with building a trio of TF, its TG, and related SNP and then test the significant coefficient difference among different levels of SNP in terms of TF and TG correlation. We applied this ceQTL model to simulated data and the lung tissue datasets from the GTEx project. The simulated data results showed that the model was robust to detect true ceQTLs at variable sample sizes and different minor allele frequencies as measured by Area Under the Curve (AUC). In normal lung tissue, a small fraction of eQTLs were found to have strong ceQTLs, i.e., eQTLs where SNP affects gene expression though TF binding. Some ceQTLs may not be detected by traditional eQTL analysis. Our tool also performed a TF binding affinity analysis to add another layer of evidence for functional interpretation. Comparisons with other similar tools were also presented. In summary, ceQTL analysis provides a more interpretable and biological insight into the mechanism of eQTL, which would help us better understand how genomic variants affect phenotypes and diseases.

## Introduction

Genome-wide association studies (GWAS) have identified an enormous number of genomic variants that are associated with various human traits or diseases [1-6]. The interpretation of variants in protein-coding regions is generally inferred from the observation that a variant changes the amino acid sequence of the protein, while those in non-coding regions, which constitute the vast majority of the variants, pose significant challenges. One of the common solutions is to conduct expression quantitative trait loci (eQTL) analysis or mapping between a genomic variant and its nearby (cis-eQTL on the same chromosome) or distant genes (trans-eQTL on a different chromosome) by linear regression [7]. Over four million genetic variants (approximately 43% of all variants with MAF ≥ 0.01) have been identified as eQTL variants, impacting about 23,000 genes in at least one of 49 tissues (referred to as cis-eVariants) from the GTEx project [8] or in the most commonly used blood samples [9].

Despite the advances, several critical questions remain. First, what is the mechanism through which an eQTL variant impacts gene expression? Second, while GWAS studies have identified tens of thousands of genomic variants associated with clinical traits or diseases, plausible explanations for most associations remain unknown, primarily due to the noncoding nature of most GWAS hits within the human genome. To address these questions, a linear regression model to test the significance of an interaction term between transcription factor (TF) and SNP effect has identified over 10K TF-eQTL interactions in about 2,000 genes from GTEx multi-tissue datasets [10]. These TF-eQTL SNPs tend to be colocalized with reported GWAS loci, which provide more evidence that the disease-associated variants affect TF binding as a mechanism; as an example from that approach, IKZF1, a TF, regulation of its TG APBB1IP through eQTL, was found to be related to various blood cell traits.

Here, we propose a new alternative model to detect ceQTLs based on Chow statistical test, which was initially proposed by Gregory Chow to test whether two linear regression models that are statistically equal [11]. It is used to detect structural breaks, where the significance of intercepts or coefficients in the models may be substantial. Nevertheless, such significance may remain undetected using a linear model with interaction terms (linear model X in short thereafter) [12]. The Chow test is widely utilized to compare parameters in two regressions but could be extended to three regressions for bi-allelic variants with three genotypes: reference homozygous, alternative allele homozygous, and heterozygous. It was tried to test if genetic variants affected patients’ Alzheimer’s disease (AD) progression based on clinical baseline variables such as cognitive and PET-based biomarkers [13]; however, to our best knowledge, this model has not been applied to eQTL analysis where the mediators can be discovered between a variant and its TGs.

## Materials and Methods

### GTEx normal lung tissue data

The pre-processed gene expression (Lung.v8.normalized_expression.bed.gz) and whole genome genotype data (GTEx_Analysis_2017-06-05_v8_WholeGenomeSeq_838Indiv_Analysis_Freeze.SHAPEIT2_phased.vcf.gz) were downloaded from the dbGAP of the GTEx project. These were the same datasets used by the GTEx project to generate eQTL results. The gene expression data was used in its original form, and the genotype data was further filtered to retain only SNPs with minor allele (MAF) > 5% on chromosomes 1-22. The meta sample data with 68 covariates were also downloaded and used in its original form. Finally, matching samples between expression, genotype, and covariates were identified, resulting in 515 samples for the final analysis.

### TF, TG and SNP trio construction

TF binding sites, TF and TG relationship mapping for human hg38 were downloaded from TKlink (https://tflink.net/, the large-scale sets, accessed April 17, 2025) [14], a database that curates TF-TG interaction information and TF binding sites in the genome from various data sources. This large scale dataset contains 1,348 TFs, 20,120 TGs, and 6,722,723 interactions. The binding site genomic coordinates were converted into bed format and then intersected with the filtered SNPs as described above. Any SNP falling into the binding regions were kept. This file was further linked to the TG of each TF using the TF and TG mapping. Only TF binding site and its TG on the same chromosome and SNPs within 10Mb of transcription start site (TSS) were retained, which led to 12,117,194 trios consisting of 530,077, 174, and 16,001 unique SNPs, TFs, and TGs, respectively.

### ceQTL calling

To perform the Chow test, for each trio, the linear regressions between TG and TF along with all 68 covariates were run on all samples (*N*) and each genotype group, respectively, to obtain the sum of squares *S, S*_*1*_, *S*_*2*_, and *S*_*3*_. Then Chow test statistic was then calculated as:

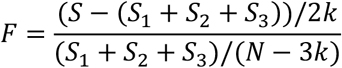

which follows the *F*-distribution with *k* the total number of parameters, including the intercept, and the *N-3k* degrees of freedom. False discovery rate (FDR) was further calculated to adjust the multiple testing for all trios tested with Q value (false discovery rate or FDR) method of R package.

### eQTL calling

To compare the ceQTL result with regular eQTL and identify those eQTLs likely modulated through TF, we performed regular eQTL analysis on all SNPs in the trios with TGs using Matrix-eQTL [15] with the default linear model as performed by GTEx.

### Linear model with TF and SNP interaction

An alternative way to detect eQTL modulated through TF is to add TF and SNP interaction in the regular eQTL model. We performed such analysis using a linear model similar to above but with additional TF and TF:SNP (TF*SNP) in the model. The analysis used the same trios as the ceQTL analysis for comparison. The interaction term p value and adjusted p value FDR were calculated to identify significant interaction, which would indicate TF regulation on TG is impacted by the underlying SNP on the TF binding site. The same FDR cut-off was used to compare results across different models.

### Motif binding affinity test

The binding affinity test was performed to investigate the effect of a mutation on TF binding. The R package, atSNP [16], a tool for annotating and predicting the effects of single nucleotide polymorphisms (SNPs) on TF binding affinity, was used. The HOCOMOCOv11 motif database [17] was also utilized to provide comprehensive information about TF binding motifs. To visualize the binding-affinity change between the alternative allele and the reference allele, MotifBreakR [18] was used to plot the motifs with surrounding sequences and the mutations.

### Data simulation

We conducted the simulation using sample sizes ranging from 100 to 400. Larger sample sizes were not included due to convergence difficulties during the simulation. The simulation was repeated 100 times for each sample size. Genotype data for 50 SNPs was initially generated using PLINK2 [19]. Different sample sizes had slightly varied allele frequency ranges to ensure at least 5 samples in each genotype group (see Table 1). The minimal and maximum allele frequencies are set to ensure that each genotype group has at least 5 samples.

**Table1.** The minimal and maximal allele frequencies used to simulate genotype for different sample sizes.

| Sample size | Minimal allele frequency | Max allele frequency |
| --- | --- | --- |
| 100 | 0.22 | 0.78 |
| 200 | 0.15 | 0.85 |
| 300 | 0.12 | 0.88 |
| 400 | 0.11 | 0.89 |

The generation of expression data was guided by the principles outlined in the dcanr package [20], a tool utilized for the simulation and analysis of differential co-expression networks. In each simulation, a total of 100 genes were generated, with 1% to 6% designated as TFs. Each TF was associated with 1 to 6 potential true positive TGs, contingent upon the degree of convergence. The correlations for these potential true positive TF-target pairs were preset, with the minimal difference in correlations across any pair of genotype groups established at 0.5. Under these parameters, initial expression values were simulated using a beta distribution, and then the values were guaranteed to fall within the 0 to 1 range.

Subsequently, the R package optimg [21], which implements the gradient descent method, was employed to optimize these expression values in alignment with the preset correlations for potential true positives. The optimization process was concluded once the system achieved a state of stability.

## Results

### ceQTL model identifies genomic variants that influence TF regulation of TG expression in lung tissue

We defined ceQTL a genomic variant that modulates the regulation of a regulator (i.e., TF) on its TG. Figure 1A illustrates one potential mechanism using TF as example about how its regulation is influenced by an underlying variant on its binding site. In essence, a variant located in the TF binding site can impact the binding affinity of the TF, thus altering the gene expression of its downstream targets. This interaction can also occur in reverse, with the variant enhancing the binding of a TF. These mechanisms can lead to different distributions in the co-expression between a TF and its TG across different genotypes of the variant (Figure 1B).

**Figure 1.**
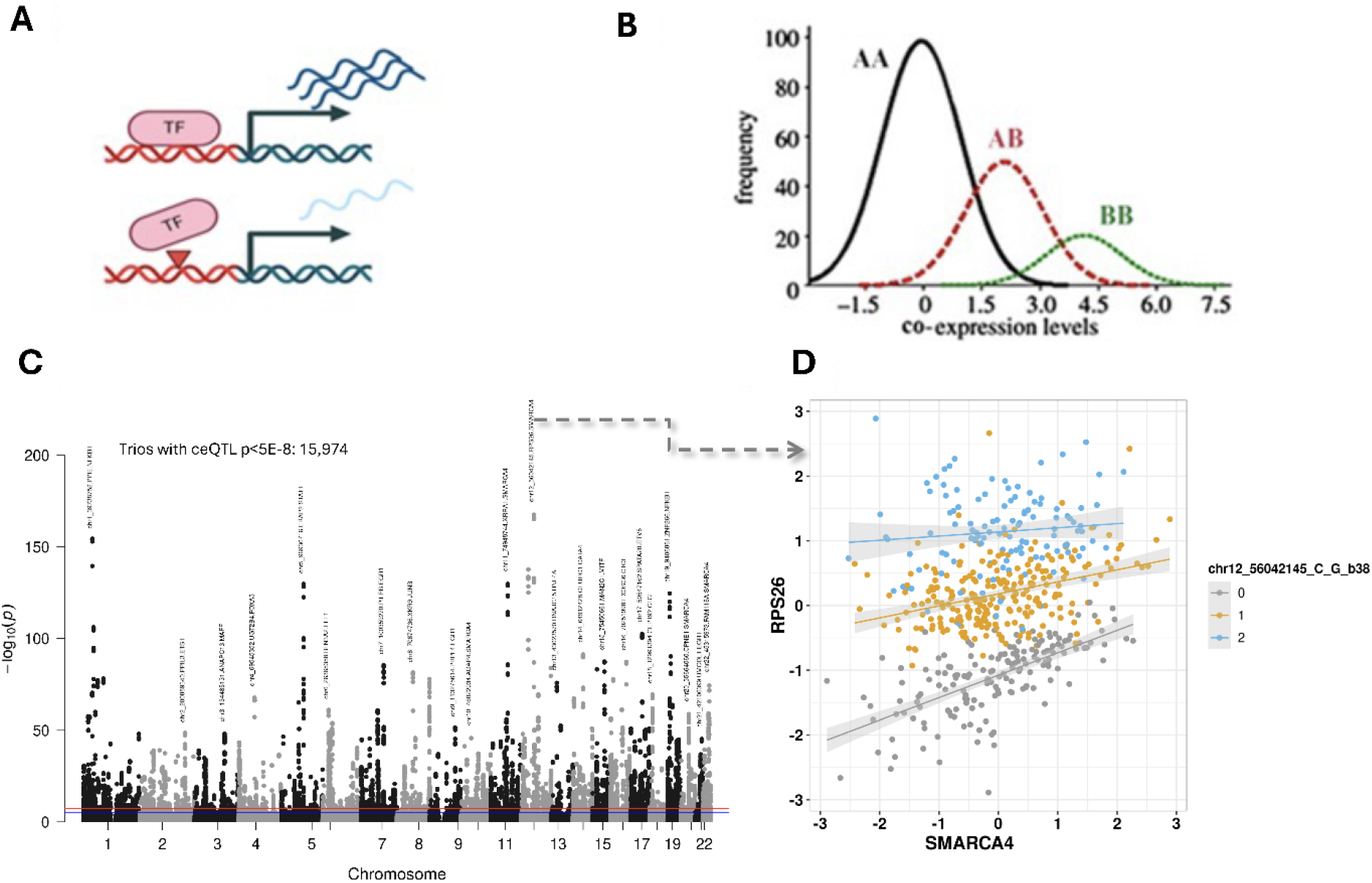
Concept and illustration of ceQTL analysis. A. The cartoon illustration of ceQTL showing the interplay among TF, a variant in its binding site, and their TG. The ceQTL models whether the variant has a significant effect on TF’s regulation of its TG expression. When there is no variant, TF up-regulates its target expression (top) while this effect is reduced when a variant is present in its binding site (bottom). B. The conceptual TG expression density plot stratified by genotype. C. The Manhattan plot of -log10 p-values for all tested ceQTL trios from lung tissues of the GTEx project. The two dashed lines marked raw p value at 1×10-5 (blue) and 1 x10-8 (red). D. A scatter plot of TF and its target expression stratified by genotypes.

We used human normal lung tissue data from the GTEx project and focused on SNPs located in the TF binding sites as curated from the TFLink database [14] (see Methods). Among the 12,117,194 mapped SNP-TF-TG trios, our ceQTL analysis identified 15,974 (0.13%) with genome-wide corrected p value less than 5E-8, and 45,800 trios (0.38%) with FDR less than 0.05 (Table 2). Figure 1C illustrates the genome-wide p value distribution (-log10 p-value) by a Manhattan plot where some top significant trios were highlighted with trio annotation. The most signfiant trio from the analysis was RPS26 (TG)-SMARCA4 (TF) -chr12_56042145_C_G_b38 (SNP) as illustrated in Figure 1D. Samples with different alleles had different levels of PRS26 expression (eQTL); at the same time, RPS26 expression was correlated with SMARCA4 expression but its correlation differed among three alleles: At homogeneous variant C RPS26 was at lower expressin and was positively correlated with SMARCA4. With one or two minor ellele G, RPS26 expression was getting higher and higher while its correaltion with SMARCA4 got reduced or dispeared (Figure 1D).

**Table 2:** Numbers of significant eQTLs and SNP-TF-TG trios by different algorithms at different significant cutoffs.

|  | eQTL | ceQTL | eQTLx (interaction) |
| --- | --- | --- | --- |
| **P<5E-8** | 69,655 | 15,974 (9,323) | 120 (108) |
| **FDR<0.05** | 230,596 | 45,800 (28,524) | 310 (273) |
| **P<0.05** | 837,548 | 612,748 (415,996) | 697,744 (656,196) |

#The number in parenthesis is for unique TG and SNP pairs

As Chow Test detects correlation structural break caused by both regression intercept and correlation coefficient, we further narrowed the significant ceQTLs to those whose TF and TG correlation p value at least or less thant 0.05. This led to 3,585 (out of 15,974 or 22.4%) trios significant at 5E-8 (Figure 2A). When a more stringent p value 5E-8 was also used for TF, only 13 trios passed the threshold involving 2 TFs (STAT1 and FOXA1), 3 TGs (TRIM69, SECTM1, and DNAH5) and 13 SNPs (Figure 2B). Two trios are plotted in Figure 2C to illustrate the signficant correlation between TF and TG and their correlations are different among different genotypes (different correlation betas). DNAH5 is very lung tissue specific and has the highest expression among all tissues profiled in GTex samples. It also has high variability among differernet individual samples (https://gtexportal.org/home/gene/DNAH5). Mutations of this gene is linked to ciliary dyskinesia [22,23]. Although TFs are generally not tissue specific, FOXA1 plays a significant role in lung airway epithelial cell functions [24].

**Figure 2.**
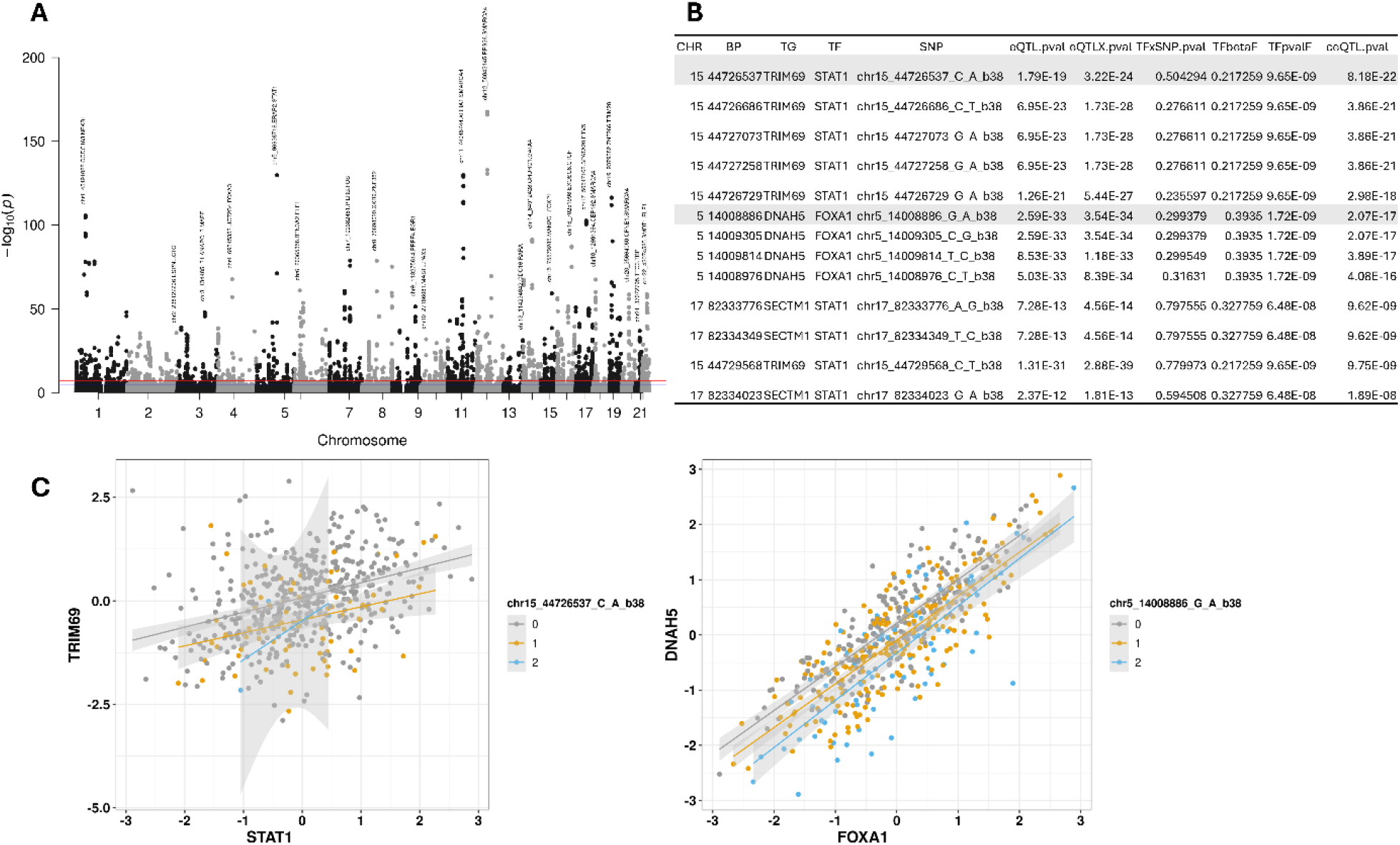
ceQTLs with ceQTL p<5E-8 and also significant TF and TG correlation in the full model (all samples regardless of genotypes). A. Manhattan plot of the tested trios with the most significant ones annotated with trio information. B. list of trios whose ceQTL p value and TF:TG correlation p value both < 5E-8. C. two of the most significant trios from 2B to illustrate TF-TG-SNP relationship.

### Comparison of ceQTLs with eQTL and linear model with interaction

The introduction of the ceQTL model extends the eQTL model by incorporating an additional layer, regulator expression of TG in the analysis, to identify those eQTLs that are mediated by TFs. When a SNP affects TF binding and subsequently alters its TG expression, it would change the correlation relationship between the TF and TG. Another approach is through detecting TF and SNP interaction in the commonly used linear model [10]. We compared these two approaches with the regular eQTL analysis.

At genome-wide significant p value less thant 5E-8, the traditinal eQTLanalysis found 69,655 SNP-TG pairs while the ceQTL identified 15,974 SNP-TG pairs (with TF as mediator) and the linear model X detected 120 TF * SNP interactions. At FDR < 0.05, the signifianct pairs were higher, with 230,596, 45,800, and 310, respectively (Table 2). Only about 10% eQTLs were found to be likely mediated by TF by ceQTL analysis; however the percentage was much lower (< 0.2%) by SNP and TF interaction analysis (Figure 3A, by p value < 5E-8). By requring TF to be significantly associated with TG expression as well, the eQTLs that were regluated by TF and impacted by SNP accounted about 2% by ceQTL analysis but the linear model X found less than 0.001% (Figure 3B).

**Figure 3.**
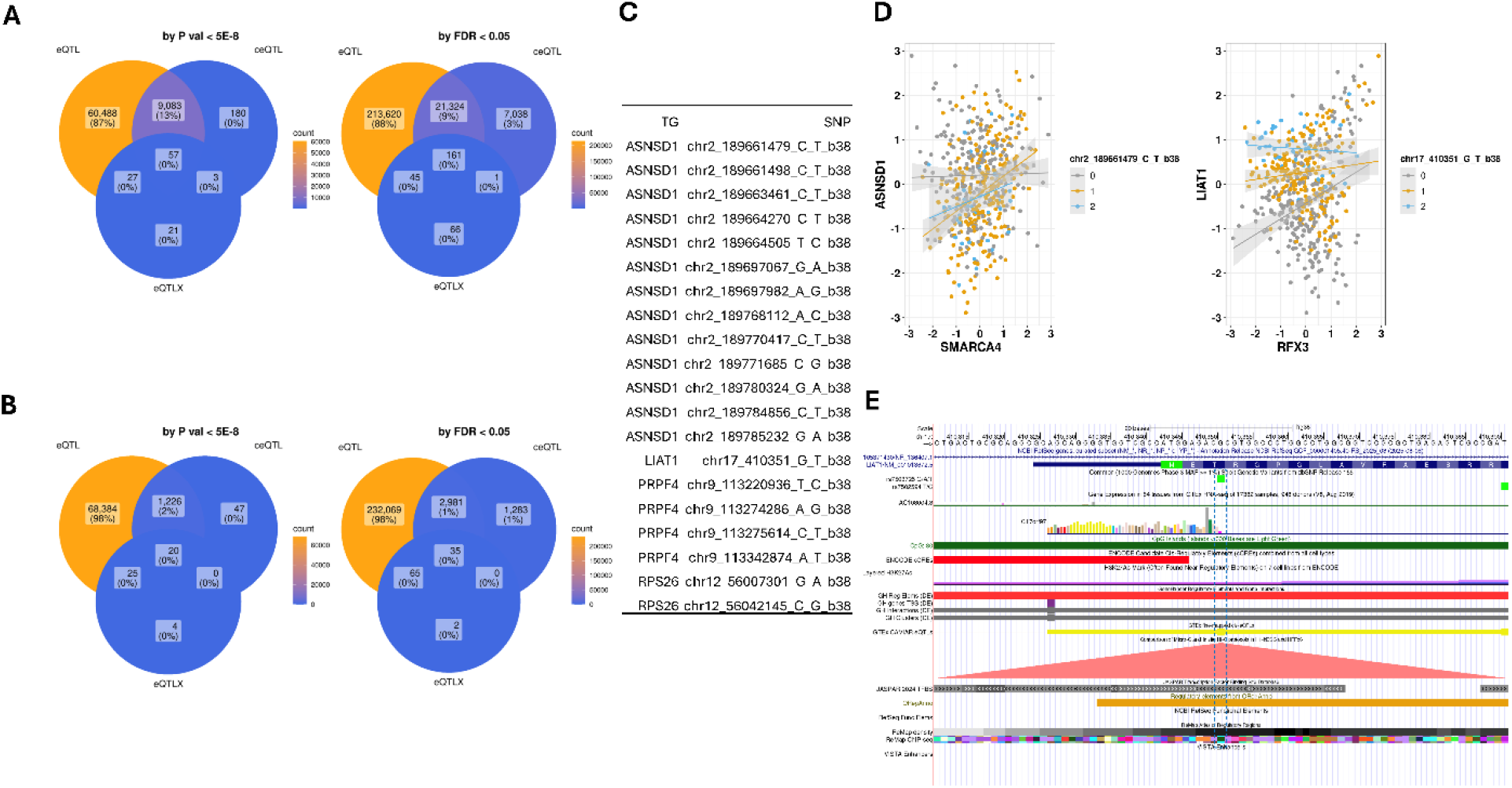
ceQTL comparison with regular eQTL and linear model X. A. Venn diagram of significant SNP-TG pairs between three methods at p value less than 5E-8 and FDR<0.05. B. ceQTLs and linear model X whose TF and/or SNP are also significantly associated with TG (p value < 0.05). C. The common TG-SNP pairs among the three methods where ceQTL and linear model X p value < 5E-8 and TF-TG correlation p value also < 5E-8. D. two top trios from 3C to show TG-TF-SNP relationship by scatter plot. E. UCSC browser view of SNP chr17_410351 (rs7503725) at the TF binding site of RFX3 where it is also at promoter region of TG LIAT1.

The three analyses found 20 SNP-TG pairs in common at the stringent p value 5E-8 (Figure 3C), with 4 unique genes, ASNSD1, LIAT1, PRPF4 and RPS26. An example plot of ASNSD1 and LIAT1 was illustrated in Figure 3D where both genes were correlated with their respetive TF (SMARCA4 and RFX3, respectively) but their relashionship was chagned by different genotypes. For example, LIAT1 was postively correlated with RFX3 at referecne allele (GG) of chr17_410351 (rs7503725) but his relationship was abolished in samples with heterozylous variant (GT) and even became negatively in homozygous variant allele (TT). The SNP sits in the promoter of LIAT1 and also the TF binding site with multipe evidences as an enhancer region (Figure 3E)

The comparison demonstrated the clear difference between the ceQTL and linear model X. Vast majoirty of eQTLs (about 87∼98% depending on filters applied) were lack of statistical evidence through TF as evidenced by either ceQTL or linear model X. ceQTL identified much more eQTLs that were likely mediated through a TF. In the more stringent results from both by P value less than 5E-8 (Figure 3B), ceQTL reported 129 SNP-TG pairs while linear model X identified 49 only, with 20 in common between the two. It is interesting to note that both ceQTL and linear model X identify small numbers of trios that were not reported by either another algorithm or regular eQTL. Careful examination of those found that some were simply below the defined significant cutoffs. For example, the trio TMEM220-FOXA1-chr17_10698004_C_A_b38 was claimed by linear model X with interaction p value 1.53E-14 while the ceQTL and eQTL p value was 2.89E-07 and 4.53E-06, respectively. On the scatter or box plot stratified by genotype, clear eQTL and SNP-TF interaction were seen (Figure 4A, B). There were some cases, howerver, where eQTL did not show any evidcne of TG expression by a SNP but ceQTL or linear model X showed a signifcant interacation between the SNP and a TF as shown in Figure 4C and D. ATP5F1B did not show any expression difference among three genotypes; however, ceQTL showed that samples with homozygous alternative allele had stronger correlation than those with homozygous reference allele or heterozygous allele. This interaction was also picked up by linear model X although its p value was just under 0.05 (Figure 4 C and D). These data suggest different algorithms can complement each other and reveal some complex or subtle relationship between SNP, TG and TF.

**Figure 4.**
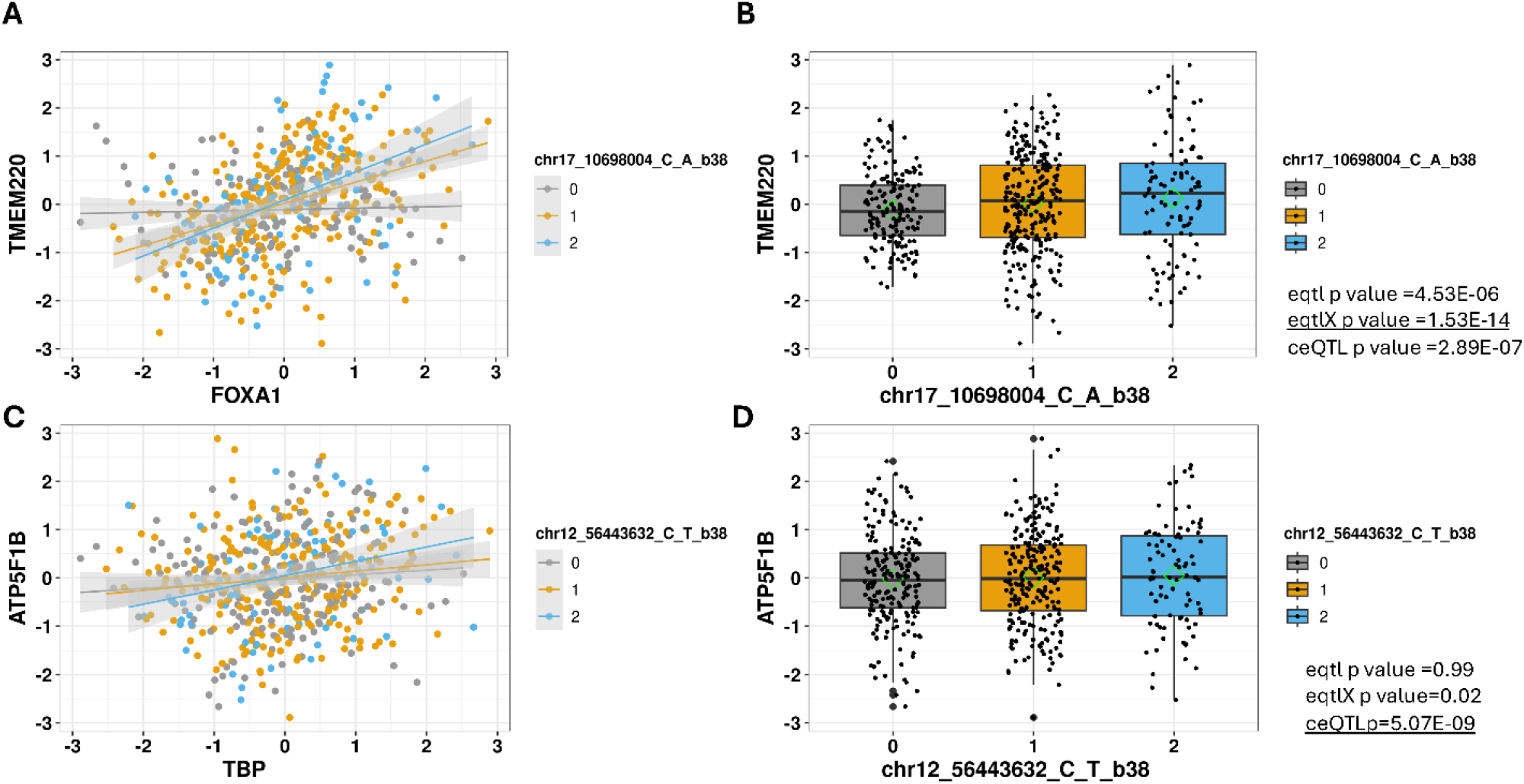
Illustration of discrepancies among ceQTL, linear model X and eQTL. A-B. significant trio claimed by linear model X but not by ceQTL and eQTL. Careful examination shows the latter are still significant but at below the set cut-off. C-D. A trio only claimed by ceQTL but not by two others.

### Case study of SNP affecting TF binding and TG expression

Among the significant ceQTLs (5E-8), we evaluated their binding affinity difference among different alleles at the TF binding site by atSNP [16] and found 70 SNP-TF combinations (44 unique SNPs and 32 unique TFs) with binding difference between reference and alternative allele as assessed by “pval_rank” less than 0.05 (49 by pval_diff, Table 3 for those significant by both). The trio ROM1:EGR1: chr11_62601964_C_G_b38 is one of the significant ones where different genotypes affected the correlation relationship between ROM1 and EGR1 (Figure 5A) and the alternative allele G reduces the binding of EGR1 about 23 fold (log likelihood difference, Figure 5B). The variant (rs12049914) sits in a CpG island, an enhancer region of ROM1, and a center of hiC interaction site (https://genome.ucsc.edu/index.html; accessed 9/30/2025). EGR1 is a very important TF in lung diseases including cancer. It is involved in cell proliferation, differentiation, apoptosis, adhesion, migration, and inflammatory responses such as chronic obstructive pulmonary disease, asthma, and pneumonia [25]. It has been also reported to be associated with lung cancer aggressiveness, PTEN status and patients’ survival [26,27]. Although ROM1 mainly is a protein in retinal outer segment, it is also reported in lung cancer as tumor suppressor [28].

**Table 3.**
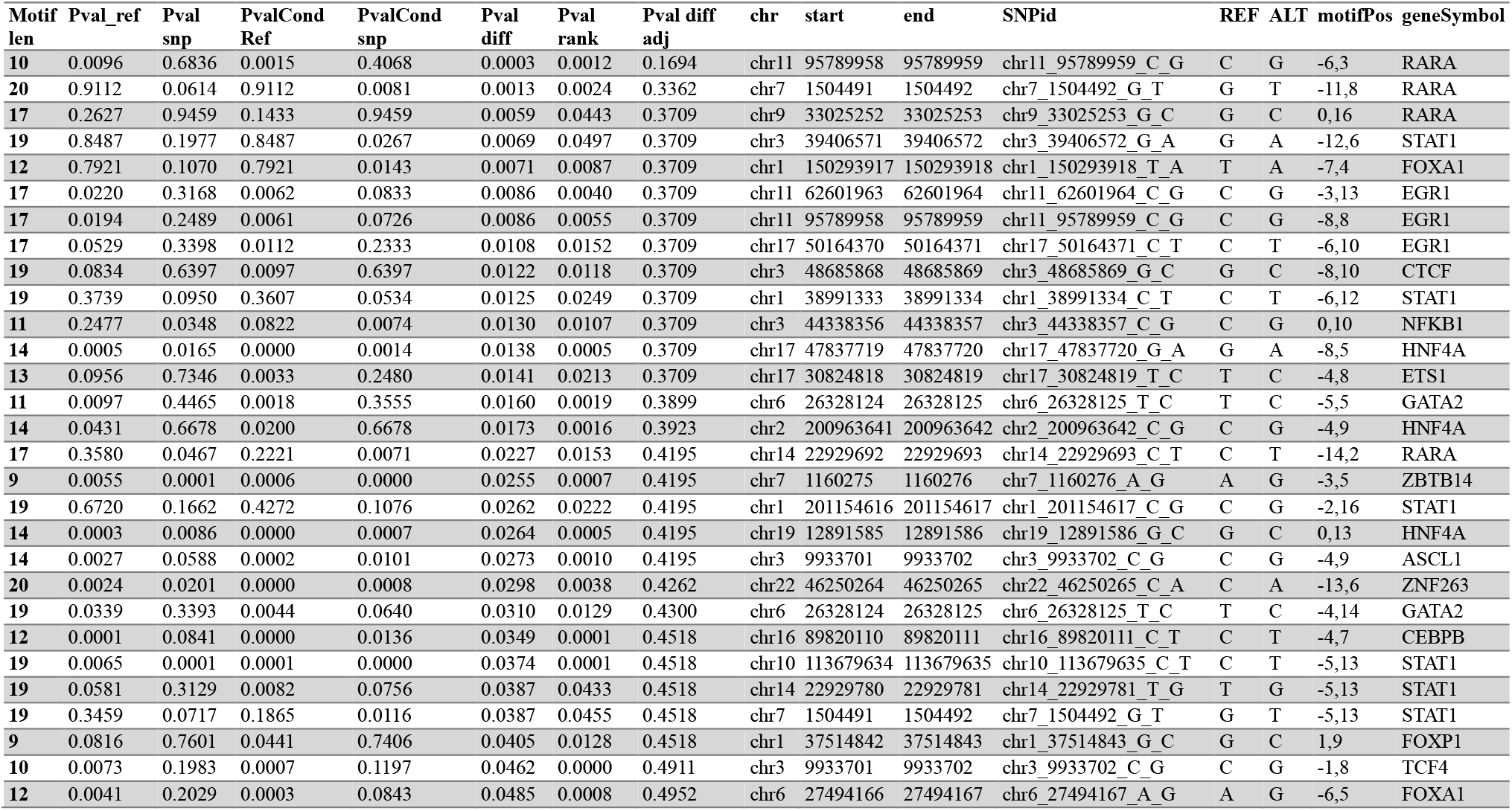
Top SNPs and binding TFs where alternative SNP affect binding affinity of TF.

**Figure 5.**
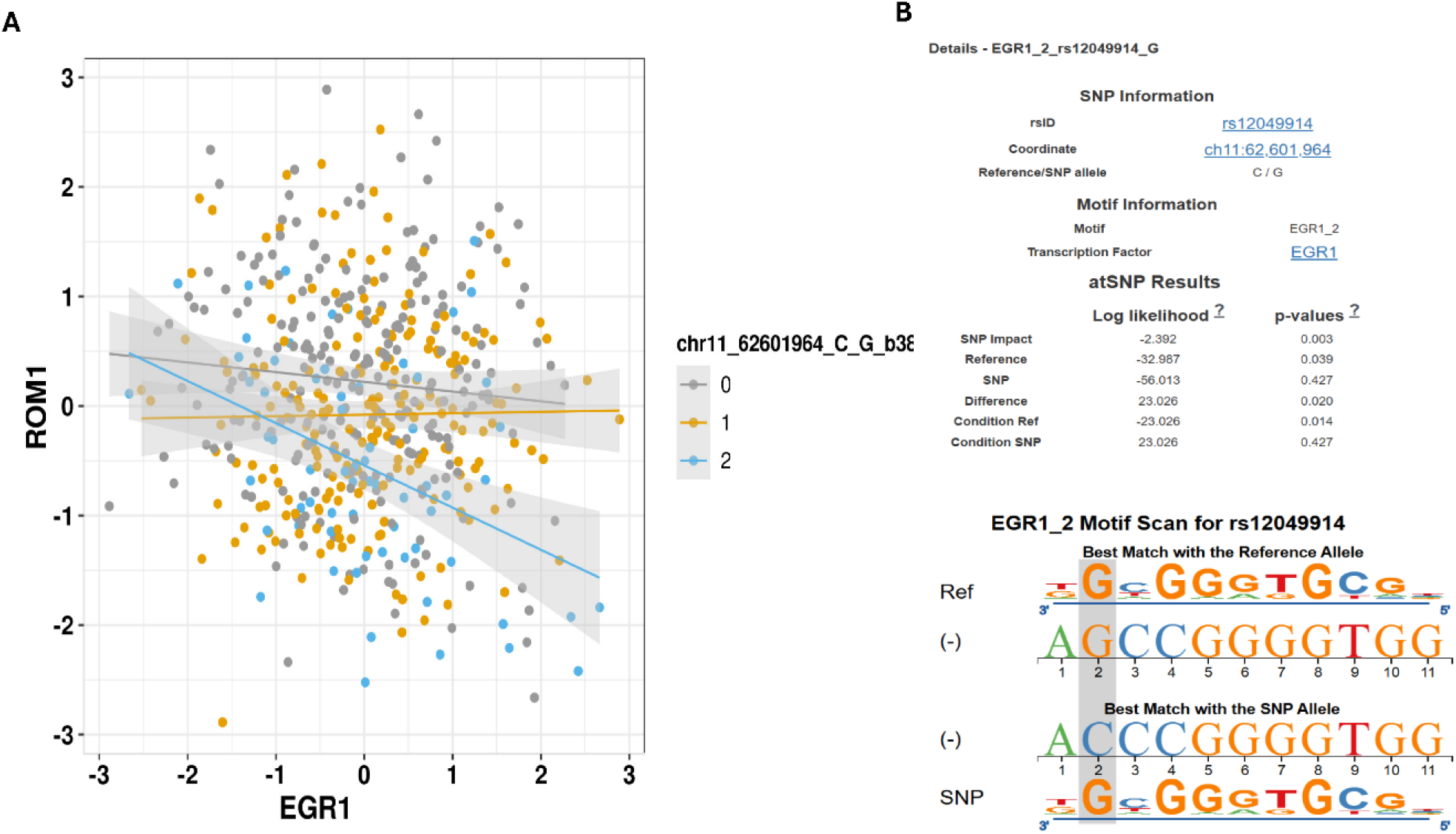
Example of a significant ceQTL trio where the TF binding site SNP changes binding affinity and affect its regulation on a TG. A. TG (ROM1) and TF (EGR1) expression scatter plot stratified by genotypes (minor allele of 0, 1, and 2 copies). Heterozygous minor allele G changes the relationship to more negative. B. atSNP evaluation of the SNP impact on TF (EGR1) binding. SNP G allele reduces the TF binding from -32.97 to -56.01 with 23 fold difference. The binding difference is statically difference with p value 0.02.

### The ceQTL model is robust with diverse sample sizes and variant allele frequencies (VAF)

To assess the robustness and reliability of the ceQTL model, a comprehensive series of 400 simulations was conducted, with sample sizes ranging from 100 to 400. Every sample size was subjected to 100 repetitions of the simulation process. The ground truth was defined by the difference in correlations of any pair of genotype groups (see details in Methods). The results, as depicted in Figure 5, provide compelling evidence of the consistent Area Under the Curve (AUC) values across the various sample sizes that were simulated. The reliability of the simulation is further demonstrated by the symmetrical trend observed when examining the minor allele frequency (MAF) between 0 and 0.5. This consistency implies that the model is robust and can effectively accommodate datasets with varying sample sizes. For a more comprehensive analysis, please refer to Figure S1A, which includes the AUCs calculated for different variant allele frequencies across various sample size groups. Furthermore, our investigation uncovered that the AUC values exhibited unwavering stability across a spectrum of variant allele frequencies, underscoring the model’s reliability across different genetic conditions. Figure S1B contains the calculated AUC values for various VAFs and sample sizes. These values provide insights into the performance of the respective models across different VAFs and sample sizes.

**Figure 5.**
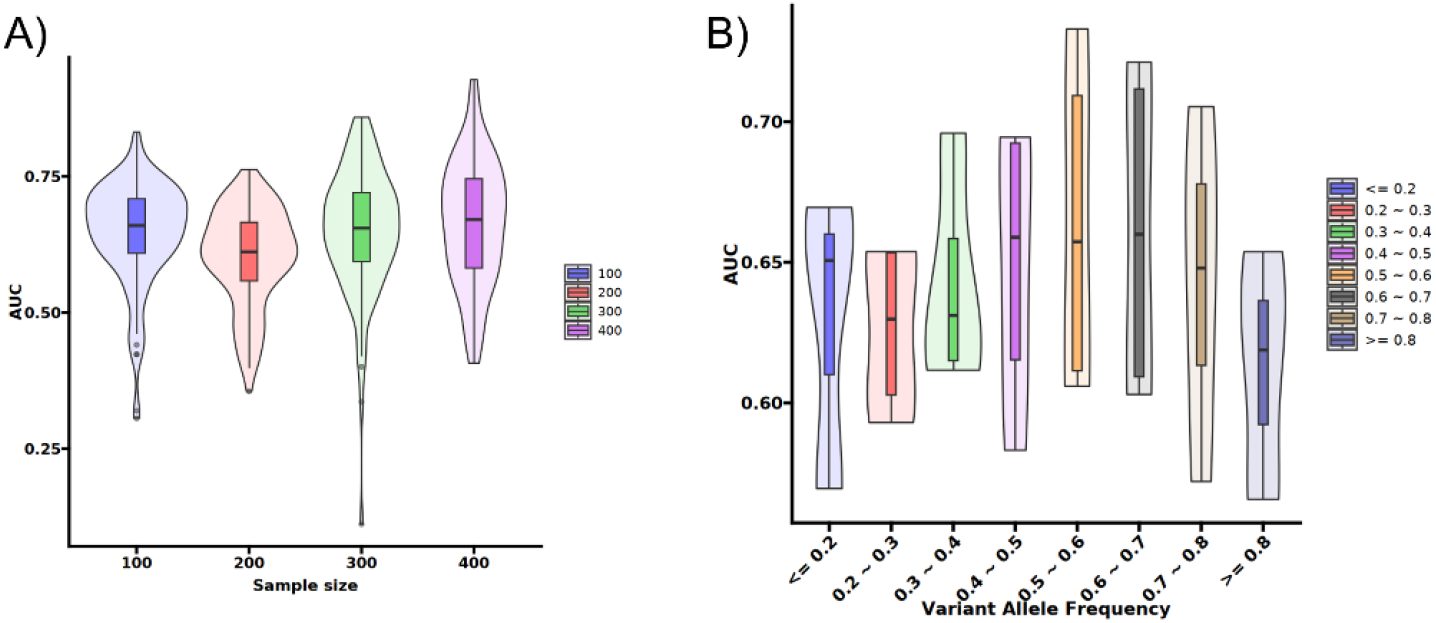
AUCs showing the robustness of the ceQTL model. A) The AUCs by different sample sizes from 100 to 400. B) The AUCs by different variant allele frequencies.

## Discussions

In this study, we have developed a new method to detect eQTL that is mediated by a TF, a step forward to interpret and explain the mechanism of gene regulation impacted by genomic variants. Our model uses the same datasets as regular eQTL analysis, but additional data preparation is needed with TF binding sites and TF and TG relationship files. We used TFLink “large-scale” datasets to be more comprehensive and inclusive although some of binding sites or interactions may have weak evidence of support. Our analysis showed that only very small fraction of eQTLs may be mediated by a TF though affecting its binding. ceQTL may also reveal some trios whose SNP and TG are not eQTL from regular eQTL analysis but by ceQTL where certain genotype changes the relationship between TF and TG. From our analysis, the SNP-TF interaction approach in the linear model appears less powerful to detect SNPs that would affect TF binding and subsequently impact its TG regulation. However, each approach has its own edges for additional trios to be discovered and combined use of these approaches may provide more comprehensive results.

ceQTL result is easier to interpret. A significant ceQTL indicates SNP changes the relationship between TF and TG significantly either positively (positive coefficient) or negatively (negative coefficient). This alone can not say for sure the change is caused by increased or decreased TF binding. The subsequent testing of binding affinity provides such evidence. Our data show that not all ceQTLs are supported by binding affinity change, suggesting other mechanisms can be involved. Additionally, when a significant ceQTL only has regression intercept difference, the SNP and TF may affect TG expression independently.

The small fraction of eQTLs that can be evidenced for TF binding interruption or enhancement by the ceQTL or linear model X may be explained that the SNP effect on TF binding may be very small yet genome wide scan requires a stringent p value cutoff to adjust multiple testing, which makes them less likely to reach the significance. The low allele frequency for a minor allele can also contribute to the underpowered statistical test, particularly when many covariates are incorporated in the model (68 variables in our analysis).

The Chow test was initially designed for two groups or levels with existing R packages (such as strucchange). This work has extended into three groups for genotype. We used TF binding site to build a trio for testing ; however, the approach is flexible and can be extended to other regions of the genome, such as open chromatin region or miRNA binding sites. Expanding the analysis to include a wider range of genomic regions, such as Hi-C [29] or 5C [30] data, particularly from the matching tissue type, will include more SNPs with TF-TG like trios in evaluation and provide a more comprehensive picture of eQTL regulation mechanisms. Similar to TF, miRNA binds to its target mRNA to perform its regulatory function and genomic variants at its binding site or itself would affect its affinity and function. A previous work [31] used a linear model to study the mechanism of eQTLs by including miRNAs as regulators.

The ceQTL model was tested using lung tissue data from the GTEx project in this work. To fully understand its robustness and applicability, it is ideal to apply the model to a broader range of datasets and tissue types in the future with more available resources. This includes other tissues from the GTEx project, data from The Cancer Genome Atlas (TCGA) [32], and other relevant repositories. Applying the model to diverse datasets will provide insights into tissue-specific regulatory mechanisms and may uncover novel ceQTLs.

### Conclusions

The ceQTL model presents a significant advancement in understanding the regulatory mechanisms linking genomic variants to gene expression. By identifying mediators such as TFs, this model provides more insights into the biological pathways affected by genetic variations. Expanding the model to include additional datasets and tissue types, refining performance metrics, and exploring alternative analytical methods will further enhance its utility and contribute to the broader field of functional genomics and disease mechanistic research.

## Author Contributions

Conceptualization, L.L., J.W., P.Y., Z.S.; Formal Analysis, P.W., and Z.S.; Data Curation, P.W., and Z.S.; Writing – Original Draft Preparation, P.W. and Z.S.; Writing – Review & Editing, all authors; Supervision, J.W., P.Y. and Z.S.; Funding Acquisition, L.L and Z.S.

## Institutional Review Board Statement

Not applicable as the data used in the study is publicly available.

## Informed Consent Statement

Not applicable (the data from public resources)

## Data Availability Statement

The GTEx gene expression and TF binding data used in the study are publicly available at GTEx (https://www.gtexportal.org/home/) and TFlink (https://tflink.net/) websites. The SNP data from GTEx has controlled access through dbGAP (accession number phs000424) and users need to apply for approval for its usage.

## Acknowledgments

The authors would like to thank The Genotype-Tissue Expression (GTEx) Project to make the data available. GTEx was supported by the Common Fund of the Office of the Director of the National Institutes of Health, and by NCI, NHGRI, NHLBI, NIDA, NIMH, and NINDS. The data used for the analyses described in this manuscript were obtained from: the GTEx Portal (https://www.gtexportal.org/home/) and/or dbGaP accession number phs000424 on 04/20/2025.

## Supplemental data

**Figure S1.**
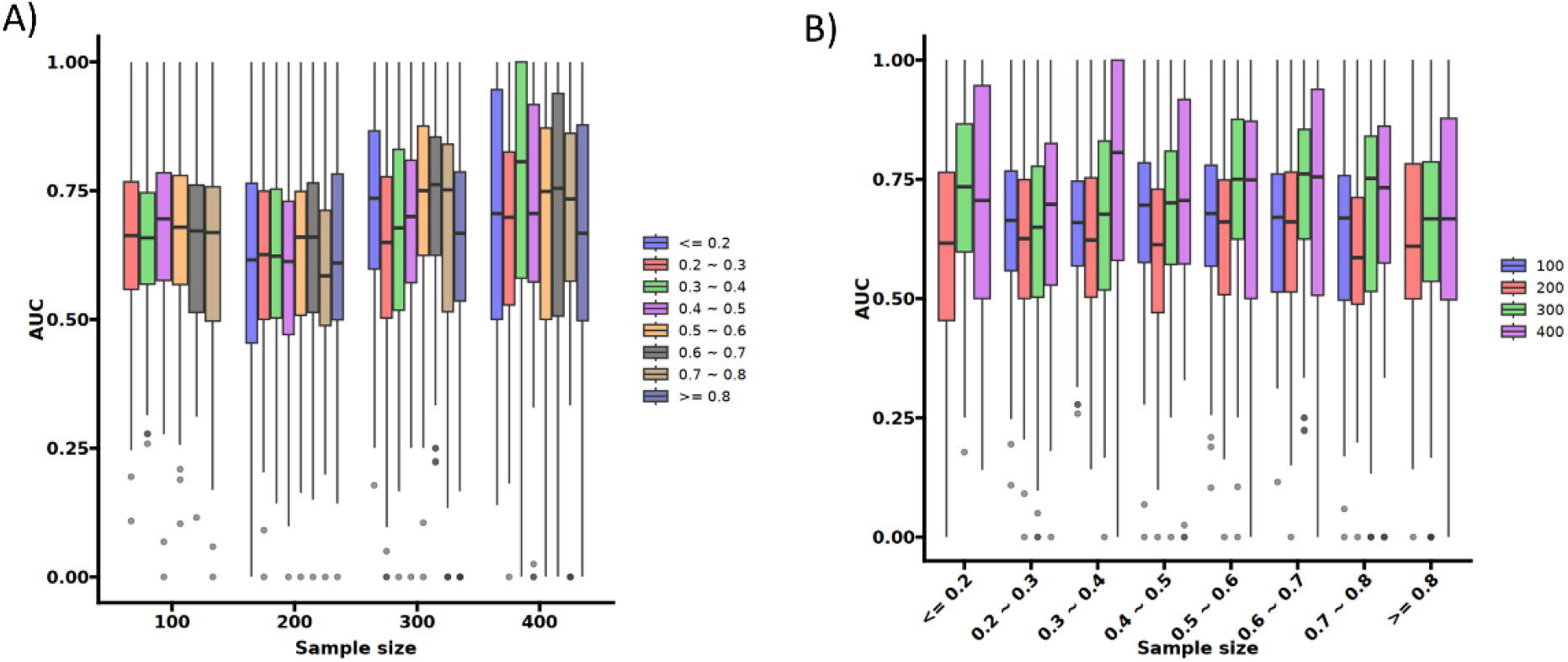
AUCs for simulated data categorized by sample sizes and variant allele frequencies. A. AUCs were calculated for simulated data using different sample sizes. At each sample size, AUCs were determined based on trios with different variant allele frequencies. B. AUCs were calculated for simulated data at different variant allele frequencies using trios of varying sample sizes.

